# Genome-wide characterization, evolutionary analysis of *WRKY* genes in *Cucurbitaceae* species and assessment of its roles in resisting to powdery mildew disease

**DOI:** 10.1101/350892

**Authors:** Zigao Jiao, Jianlei Sun, Chongqi Wang, Yumei Dong, Shouhua Xiao, Xuli Gao, Qiwei Cao, Libin Li, Wendong Li, Chao Gao

**Affiliations:** Shandong Key Laboratory of Greenhouse Vegetable Biology, Institute of Vegetables and Flowers, Shandong Academy of Agricultural Sciences, Jinan 250100, China

**Author notes:** These authors contributed equally to this work (Zigao Jiao, Jianlei Sun). Corresponding author; E-mail: < >.

**Keywords:** ***cucurbitaceae***, **WRKY**, **phylogenetic relationship**, **evolution**, **powdery mildew disease**

## Abstract

The WRKY proteins constitute a large family of transcription factors that have been known to play a wide range of regulatory roles in multiple biological processes. Over the past few years, many reports have focused on analysis of evolution and biological function of *WRKY* genes at the whole genome level in different plant species. However, little information is known about *WRKY* genes in melon (*Cucumis melo* L.). In the present study, a total of 56 putative *WRKY* genes were identified in melon, which were randomly distributed on their respective chromosomes. A multiple sequence alignment and phylogenetic analysis using melon, cucumber and watermelon predicted WRKY domains indicated that melon WRKY proteins could be classified into three main groups (I-III). Our analysis indicated that no recent duplication events of *WRKY* genes were detected in melon, and strong purifying selection was observed among the 85 orthologous pairs of *Cucurbitaceae* species. Expression profiles of *CmWRKY* derived from RNA-seq data and quantitative RT-PCR (qRT-PCR) analyses showed distinct expression patterns in various tissues, and the expression of 16 *CmWRKY* were altered following powdery mildew infection in melon. Besides, we also found that a total of 24 *WRKY* genes were co-expressed with 11 *VQ* family genes in melon. Our comparative genomic analysis provides a foundation for future functional dissection and understanding the evolution of *WRKY* genes in *cucurbitaceae* species, and will promote powdery mildew resistance study in melon.

## Introduction

WRKY proteins are widely distributed in all plants and comprise one of the largest transcription factor families. Over the past decades, these proteins have been found to play an increasing number of functions in a wide range of physiological and biochemical processes [1–5]. The WRKY family proteins are defined by a highly conserved domain with approximately 60 amino acid residues in length, which contains one or two highly conserved short peptide WRKYGQK as well as a conserved C2H2- or C2HC-type zinc finger motif [6–8]. The conserved short peptide also consists of various forms, such as WRKYGKK, WRKYDQK, and WRKYDHK. Based on the number of WRKY domains and the type of zinc finger motif, the WRKY family proteins can be classified into three groups [7, 8]. Group I contains two WRKY motifs and a C2H2 type zinc finger motif (C-X4-5-C-X22-23-H-X1-H). Group II contains one WRKY motif and a C2H2 type zinc finger motif. Group III also contains only one WRKY motif but has a C2HC type zinc finger motif (C-X7-C-X23-H-X1-C). WRKY transcription factors in group II can be further divided into five subgroups (IIa, IIb, IIc, IId, and IIe) according to their phylogenetic relationship [6].

With the release of an increasing number of genome sequences, more and more *WRKY* genes have been identified in a wide range of plant species [9–14], suggesting that WRKY proteins have evolved diverse functions with different biochemical properties. To date, many plant WRKY proteins have also been functionally studied in detail, suggesting that they are implicated to modulate seed development, flowering, fruit ripening, senescence and various metabolic processes by binding to W-box element ((C/T)TGAC(T/C)) in the promoter of downstream target genes [15–19]. For example, a WRKY protein is strongly expressed in the endothelium of developing seeds, and plays important role during seed coat development in *Arabidopsis* [20]. In soybean, differential expression of *GsWRKY15a* between wild and cultivated soybean pods was correlated with different seed size [19]. Besides, VvWRKY26, a TTG2-like homolog protein, controls vacuolar acidification, transport and flavonoid biosynthesis in grapevine, and plays important regulatory roles in fleshy fruit development [17].

Recent studies revealed that a large part of WRKY proteins were also involved in response to biotic (bacterial, fungal and viral pathogens) defense and abiotic stress (salinity, drought, heat, cold and osmotic stress) [21]. In many plants, expression of *WRKY* family genes was significantly up- or down-regulated in response to drought and salt stress. Furthermore, overexpression of *TaWRKY2* and *TaWRKY19* confer tolerance to salt, drought, and cold stresses in transgenic *Arabidopsis* plants and overexpression of *TaWRKY44* gene in tobacco improved osmotic stress tolerance [22, 23]. Notably, increasing reports showed that the regulatory roles of WRKY proteins in plant defense response are closely associated with hormone-mediated signal pathways such as salicylic acid (SA) signaling pathway, abscisic acid (ABA) signaling pathway, jasmonic acid (JA) signaling pathway and so on [24–26]. For instance, various phytohormone treatments significantly altered the expression patterns of 54 *WRKY* genes in rice [27]. AtWRKY50 and AtWRKY51 work as positive regulators in the SA signaling pathway but as negative regulators in JA signaling pathway [25].

Melon (*Cucumis melo* L.), watermelon (*Citrullus lanatus* L.) and cucumber (*Cucumis sativus* L.) are economically important fruit crop that belong to *cucurbitaceae* family. Recently, the genome sequences of melon, watermelon and cucumber have been released, which provides an important reference for genome-wide analysis of gene families and compare evolutionary relationship among their homologs [28–30]. In the previous study, *WRKY* family genes in cucumber and watermelon have been respectively identified and characterized. Meanwhile, their structure, chromosome distribution, phylogeny, conserved motifs as well as expression patterns under various abiotic stresses were also analyzed, which provide important information for the evolutionary analysis and functional characterization of the *WRKY* gene family among cucumber, watermelon and other species [11, 14]. However, the basic knowledge of the melon *WRKY* family genes as well as the evolutionary relationship, functional conservation and diversification among *WRKY* family genes in *cucurbitaceae* species have still not been reported.

Thus, our aim was mainly focus on the identification and characterization of *WRKY* family genes in melon and analysis the evolutionary relationships of *WRKY* family genes among *cucurbitaceae* species. In the present study, we identified a total of 56 proteins with complete WRKY domains in melon using a hidden Markov model (HMM) that allows for the detection of the WRKY domain across highly divergent sequence. Multiple sequence alignments, phylogenetic relationships, chromosome distributions, gene duplication, syntenic relationship and selection pressure analysis of *WRKY* orthologous pairs from *cucurbitaceae* species were also performed. In addition, this study also determined the expression patterns of *CmWRKY* genes in 29 different tissues and measured their abundance under powdery mildew fungus infection using quantitative RT-PCR (qRT-PCR). Furthermore, we also performed the correlation analysis between *CmWRKY*s and *CmVQ*s expression using the data from RNA-seq. Our results will undoubtedly provide a foundation for further evolution and functional studies of *cucurbitaceae* WRKY family genes.

## Materials and Methods

### Plant materials and powdery mildew fungus inoculation

Powdery mildew fungus was collected from cultivated melon grown on the experimental farm of Shandong Academy of Agricultural Sciences with normal day/night period. Cultivated melon (B29) were grown in the greenhouse with a photoperiod of 16/8 h (day/night) and a temperature of 22°C/12°C (day period/night period). Plants with two or three true leaves were inoculated by powdery mildew fungus with a concentration of 1× 10^6^/mL as previously described [31]. Leaves were harvested at 0, 24, 72, 168 h post inoculation, and immediately frozen in liquid nitrogen and stored at −80°C for the following RNA extraction. Three biological replicates were prepared for each sample.

### Identification of candidate *WRKY* gene family members in melon

The Hidden Markov Model (HMM) of the WRKY domain (PF03106) was downloaded from the Pfam protein family database (http://pfam.sanger.ac.uk/) and used to identify putative WRKY proteins from the melon genome sequence (http://cucurbitgenomics.org/) using the HMM search as well as BLASTP search program with default parameters (HMMER 3.0; http://hmmer.janelia.org/). All non-redundant protein sequences encoding complete WRKY domains were selected as putative WRKY proteins and confirmed using the SMART software program (http://smart.embl-heidelberg.de/).

### Multiple sequence alignment and phylogenetic analysis

Multiple sequence alignment was determined using Clustal Omega online software (http://www.ebi.ac.uk/Tools/msa/clustalw2/) with default settings using amino acid sequences of all WRKY proteins as input. Subsequently, MEGA7.0 software was used for phylogenetic analysis using the neighbor-joining method with 1000 replicates of bootstrap based on the alignment results. The phylogenetic tree showed only branches with a bootstrap consensus > 50. Based on the multiple sequence alignment and the previously reported classification of *CsWRKY* and *ClWRKY* genes, the *CmWRKY* genes were assigned to different groups and subgroups.

### Chromosomal location, gene duplication and selection pressure analysis

The chromosomal location information of all melon *WRKY* genes was obtained from cucurbit genomics database (http://cucurbitgenomics.org/). The map was generated using MapInspect software (http://mapinspect.software.informer.com/). To identify gene duplication events, the nucleotide sequence within melon were respectively aligned. To detect gene duplication events, every WRKY nucleotide sequence were aligned against the other WRKY nucleotide sequence in melon, respectively. Genes which were found within the 5-Mb regions with 80% or higher nucleotide sequence similarity and e-value threshold of 1e^−10^ were considered as tandemly duplicated genes, and the ones separated by >5 Mb distance were identified as segmentally duplicated genes [32]. Non-synonymous (Ka) and synonymous (Ks) substitution of each orthologous gene pair were calculated by PAL2NAL program [33], which is based on codon model program in PAML [34]. Generally, Ka/Ks=1, >1, and <1 indicated neutral, positive, and purifying selection, respectively.

### RNA-seq based expression analysis and correlation calculation

The normalized expression levels of melon *WRKY* and *VQ* genes in different tissues as well as different developmental stages based on RNA-seq data were obtained from the Melonet DB for functional genomics study of muskmelon (http://melonet-db.agbi.tsukuba.ac.jp/cgi-bin/top.cgi). Gene expression data are presented as log_2_ (RPKM value+1) to reveal difference in expression levels among different tissues. To visualize the expression patterns of the *WRKY* genes in different melon organs, a heat map was created using R project (http://www.r-project.org/). The co-expression correlation analysis between *WRKY* genes and all *VQ* genes was performed using R programming language (http://www.r-project.org/). The co-expression network was constructed using genes with r > 0.6 and a FDR < 0.05.

### RNA isolation and qRT-PCR analysis

Total RNA samples were extracted from leaves using the Trizol reagent according to the manufacturer’s instruction (Invitrogen, CA, USA). Reverse transcription reactions were performed at 42°C for 1 h and were terminated at 85°C for 5 min with 20 μl system contained 1×reverse transcriptase buffer, 50 nM Olig(dT) primer, 0.25 mM each of dNTPs, 50 units reverse transcriptase, 4 units RNase inhibitor and 2 μg DNase I-treated total RNA. *CmActin* was used as the internal control. The qRT-PCR program was set as follows: 95°C for 5 min, then followed by 40 cycles of 95°C for 15 s and 60°C for 1 min. The 2^−ΔΔCt^ method was used to calculate the relative expression levels and t-test was used to access whether the results were statistically different (*P < 0.05). Primers used for qRT-PCR experiments were listed in S1 Table.

## Results

### Identification of WRKY family genes in melon

To identify WRKY family genes in melon, Hidden Markov Model (HMM) and BLASTP searches were performed against reference genomes of melon using the consensus sequence of the WRKY domain. As a result, a total of 56 proteins with complete WRKY domains were identified in melon, which were termed as *CmWRKY1* to *CmWRKY56*. The numbers of *WRKY* family members was approximately equal with other *cucurbitaceae* species cucumber and watermelon but less than that in model plants *Arabidopsis* (74 members) and in rice (109 members). The nucleotide and amino acid sequences of all identified WRKY families are presented in S2 Table. As shown in S2 Table, the lengths of the putative CmWRKY proteins ranged from 111 to 768 amino acids, with an average of 342 amino acids. Based on the number of WRKY domain and the type of zinc finger motif, CmWRKY proteins could be classified into three groups, group I (11 sequences), group II (40 sequences) and group III (5 sequences) based on the number of WRKY domain and the type of zinc finger motif (S3 Table).

### Multiple sequence alignment and phylogenetic analysis of WRKY domains

The WRKY domain consists of approximately 60 amino acid residues and contains one or two
highly conserved short peptide WRKYGQK as well as a conserved C2H2- or C2HC-type zinc finger motif.^7,8^ The conserved short peptide WRKYGQK is considered to be important for recognizing and binding to W-box elements. A multiple sequence alignment of the core WRKY domain in melon were performed and shown in Fig 1. WRKYGQK sequences represented the major variant in 54 melon WRKY proteins. WRKYGKK sequence was observed only in two WRKY proteins (CmWRKY16 and CmWRKY48) that belong to group IIa.

**Figure 1.**
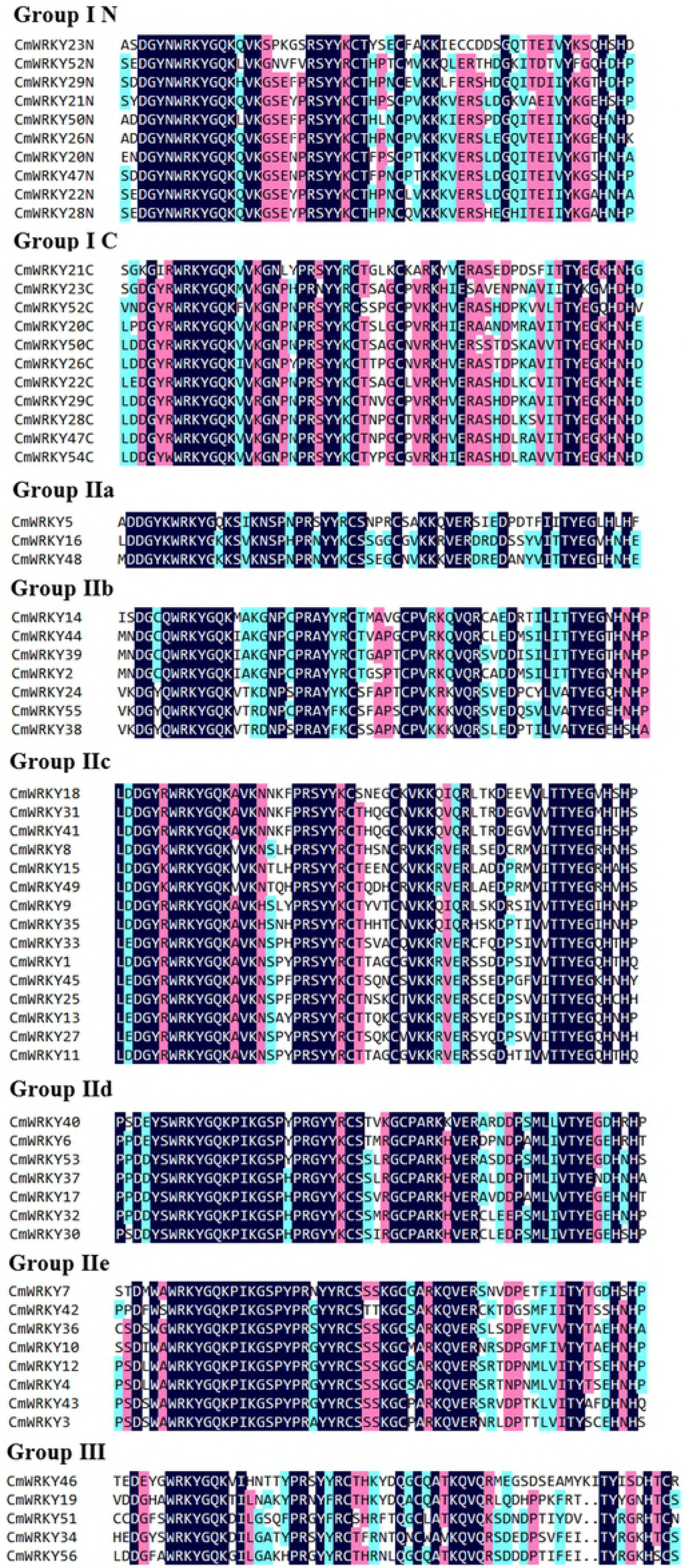
Multiple sequence alignment of 56 conserved WRKY domains in melon. Highly conserved amino acids in WRKY domain are shown in black.

To identify the evolutionary relationships of the WRKY proteins among three *cucurbitaceae* species (melon, cucumber, and watermelon), a neighbor-joining (NJ) phylogenetic tree was generated using multiple sequence alignments of all the conserved WRKY domains with a bootstrap analysis (1,000 replicates). Based the classification of WRKY domains in cucumber and watermelon, all melon *WRKY* genes were classified into three main groups, with five subgroups in group II. As shown in Fig 2, group I contained 11 melon WRKYs, 11 watermelon WRKYs and 12 cucumber WRKYs. While, group III contained 5 melon WRKYs, 8 watermelon WRKYs and 6 cucumber WRKYs. Besides, 40 CmWRKY proteins in group II could be classified into five subgroups based on the phylogenetic trees, IIa (3 sequences), IIb (7 sequences), IIc (15 sequences), IId (7 sequences), and IIe (8 sequences). Meanwhile, 34 ClWRKY proteins in group II could be classified into IIa (2 sequences), IIb (8 sequences), IIc (11 sequences), IId (6 sequences) and IIe (7 sequences), and 38 ClWRKY proteins in group II could be classified into IIa (5 sequences), IIb (7 sequences), IIc (12 sequences), IId (7 sequences) and IIe (7 sequences). Detailed information about the classification of the *WRKY* genes, as well as the sequences of conserved WRKY domains and zinc-finger motifs in each gene can be found in S3 Table.

**Figure 2.**
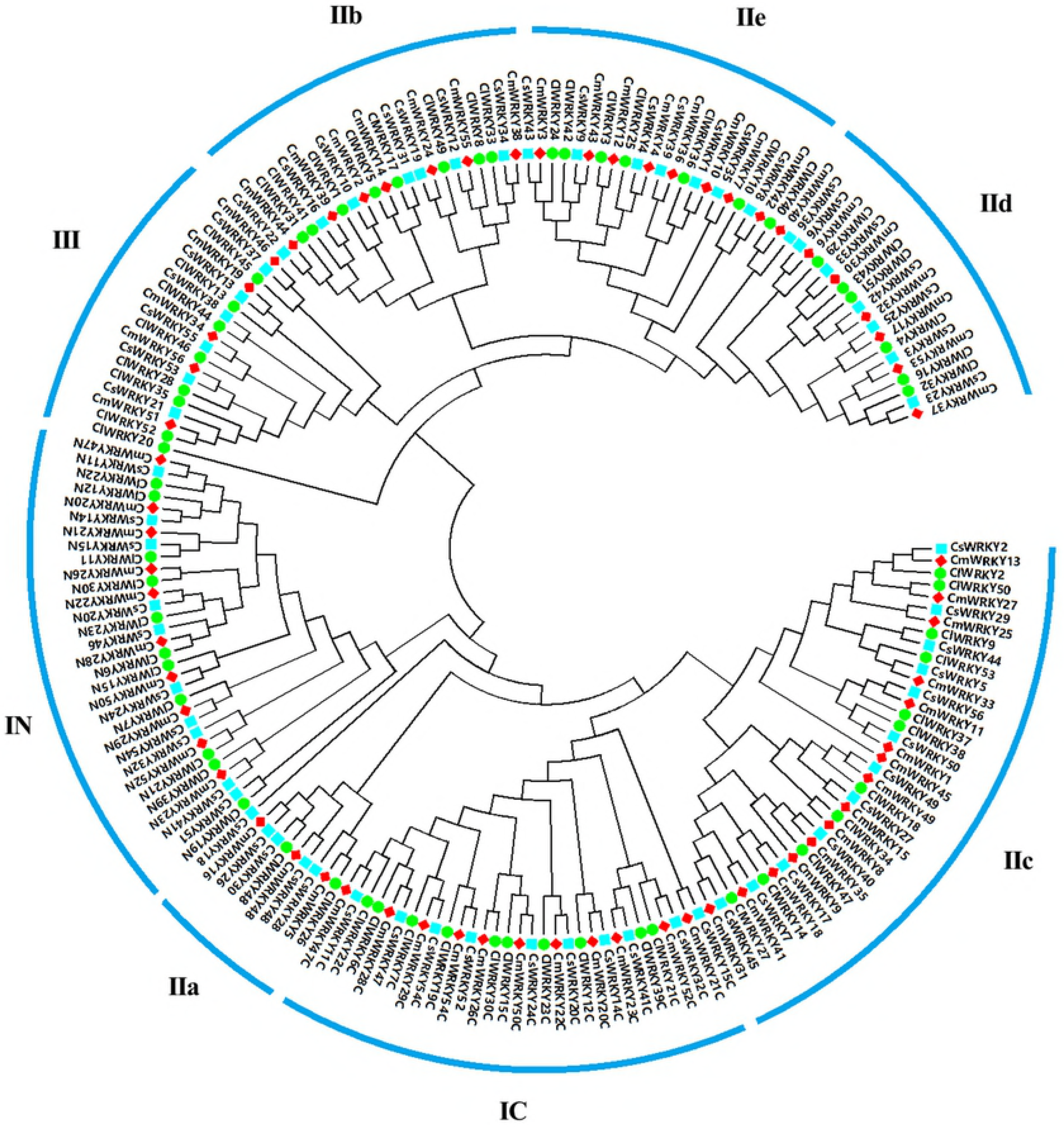
Phylogenetic relationships of WRKY domains from melon (red square), cucumber (blue square) and watermelon (green circle). The domains were clustered into three major groups I, II, III, and five subgroups (a, b, c, d, and e) in group II.

### Genome-wide distribution and duplication of *WRKY* family genes

Fig 3 showed the distribution of the *WRKY* genes on melon chromosomes. As shown in the figures, the WRKY genes were unevenly distributed throughout all chromosomes, and the number on each chromosome was not related to its length. In melon, chromosome 6 contained the largest number (9) of CmWRKY genes. To identify the contribution of segmental and tandem duplications in genome-wide expansion of WRKY family in the *cucurbitaceae* genomes, *WRKY* genes which were found within the 5-Mb regions with 80% and higher nucleotide similarity with e-value threshold of 1e^−10^ were considered as tandemly duplicated genes, and the ones separated by >5 Mb distance were identified as segmentally duplicated genes. The results showed that the highest levels of nucleotide identity were the pairs *CmWRKY4-CmWRKY6* (75.6%), *ClWRKY4-ClWRKY6* (76.8%) and *CsWRKY4-CsWRKY6* (74.2%) for melon, watermelon and cucumber, respectively. The result did not provide clear evidence of recent gene duplication events including tandem duplication and segmental duplication for each of the three species, as the nucleotide identity of *WRKY* pairs were lower than 80%, which revealed that duplication event plays an insignificant role in *cucurbitaceae WRKY* genes evolution. Indeed, Ling has discovered that no gene duplication events occurred in *CsWRKY* gene evolution because of no paralogs of cucumber can be detected through phylogenetic and nucleotide identity analysis.

**Figure 3.**
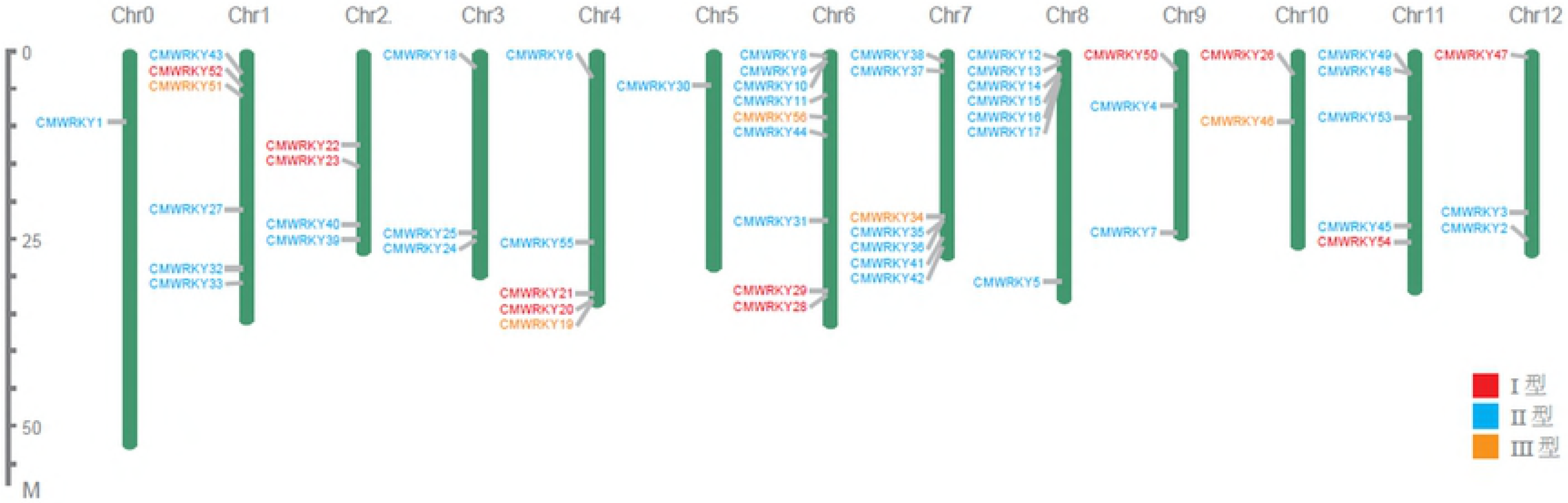
Chromosomal location of *WRKY* family genes in melon. The size of a chromosome is indicated by its relative length. The chromosome numbers were shown at the top of each chromosome.

### Orthologous gene identification and selection pressure analysis of *WRKY* orthologous genes among *cucurbitaceae* species

In our study, we further identified 44 orthologous pairs between melon and watermelon, 43 orthologous pairs between melon and cucumber, and 40 orthologous pairs between watermelon and cucumber according to the phylogenetic and homogeneity analysis (S4 Table). The highest and lowest amino acid identity between melon and cucumber were the pairs *CmWRKY50-CsWRKY24* (98.62%) and *CmWRKY7-CsWRKY35* (74.21%) with an average sequences identity of 93.00 %. The highest and lowest protein sequence identity between melon and watermelon were the pairs *CmWRKY53-ClWRKY16* (95.92%) and *CmWRKY5-ClWRKY26* (65.51%) with an average sequences identity of 83.57%. The highest and lowest amino acid identity between watermelon and cucumber were the pairs *CsWRKY24-ClWRKY15* (95.65%) and *CsWRKY37-ClWRKY45* (61.67%) with an average sequences identity of 83.93%. The chromosomal location and syntenic relationship of orthologous gene pairs were shown in Fig 4. Physical mapping revealed that most WRKY genes (98%) in *cucurbitaceae* species were not located in the corresponding chromosomes of melon, watermelon and cucumber, suggesting the occurrence of large chromosome rearrangement in the *cucurbitaceae* genomes. Furthermore, the dissimilarity level between the non-synonymous substitution (dN) and synonymous substitution (dS) values was used to infer the direction and magnitude of natural selection acting on *WRKY* orthologous gene pairs in melon, watermelon and cucumber. The results showed that the *WRKY* orthologous gene pairs in *cucurbitaceae* species underwent strong purifying pressure during evolution (Table 1).

**Table 1.**
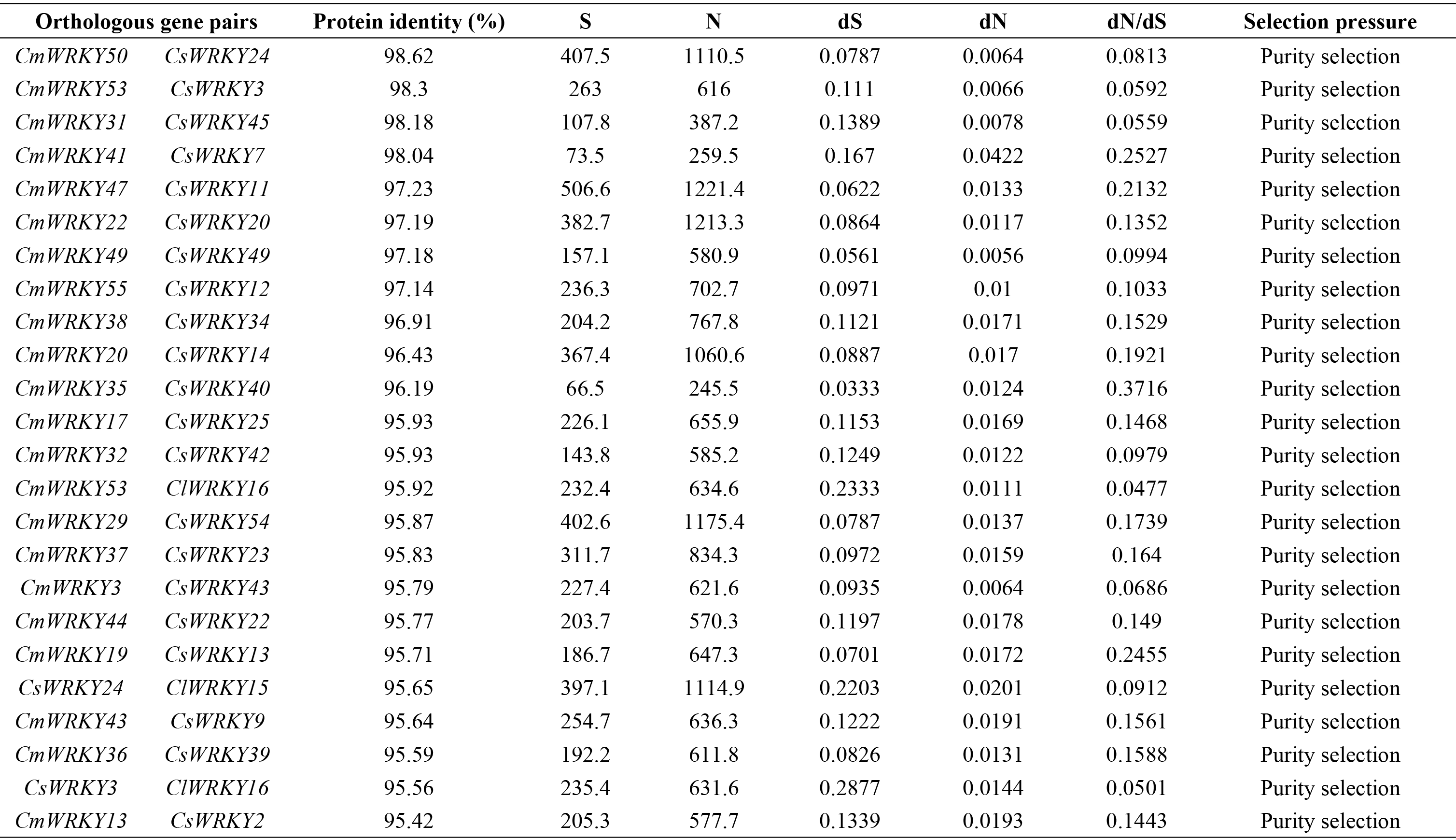
Ka/Ks calculation of each orthologous pairs among three *cucurbitaceae* species.

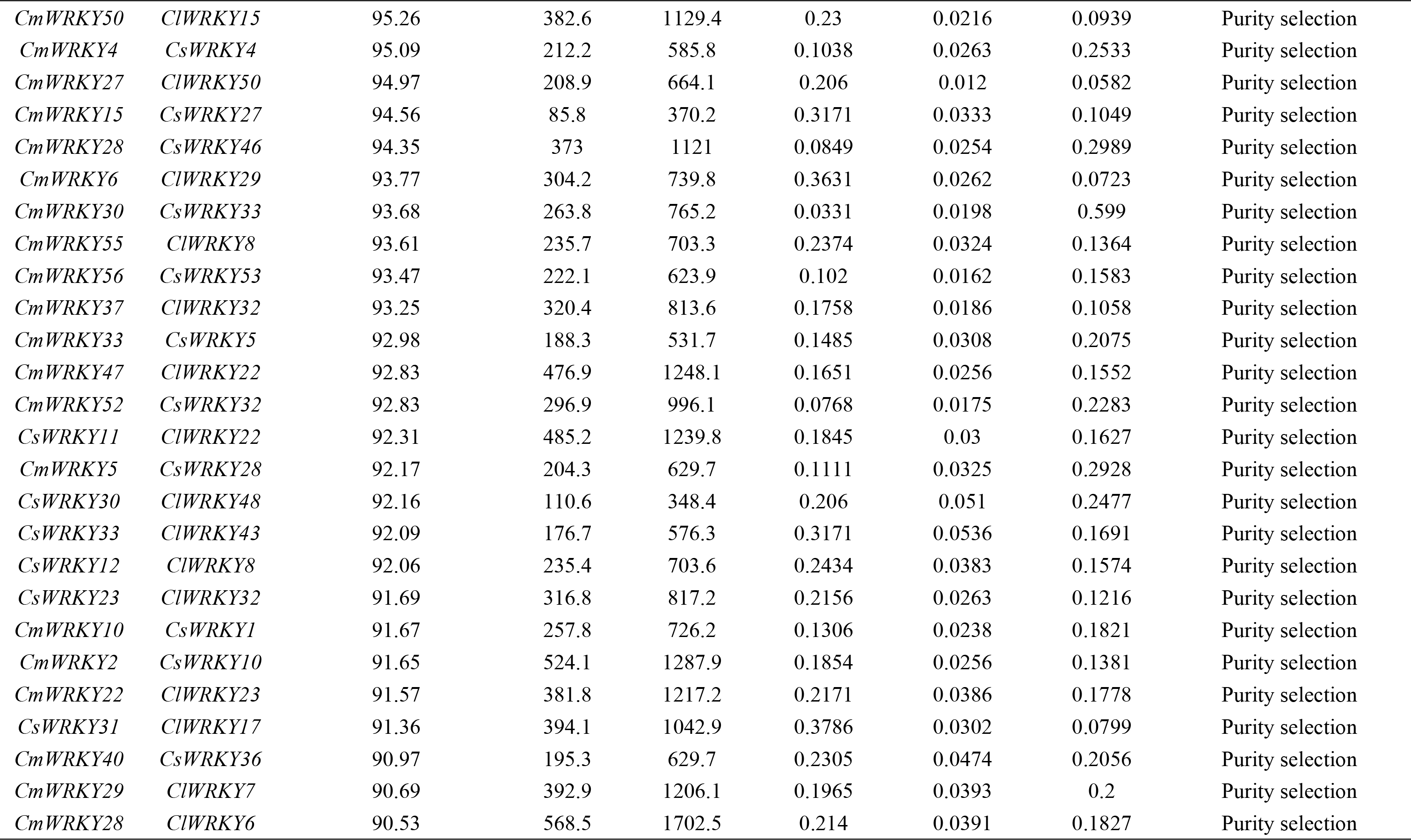

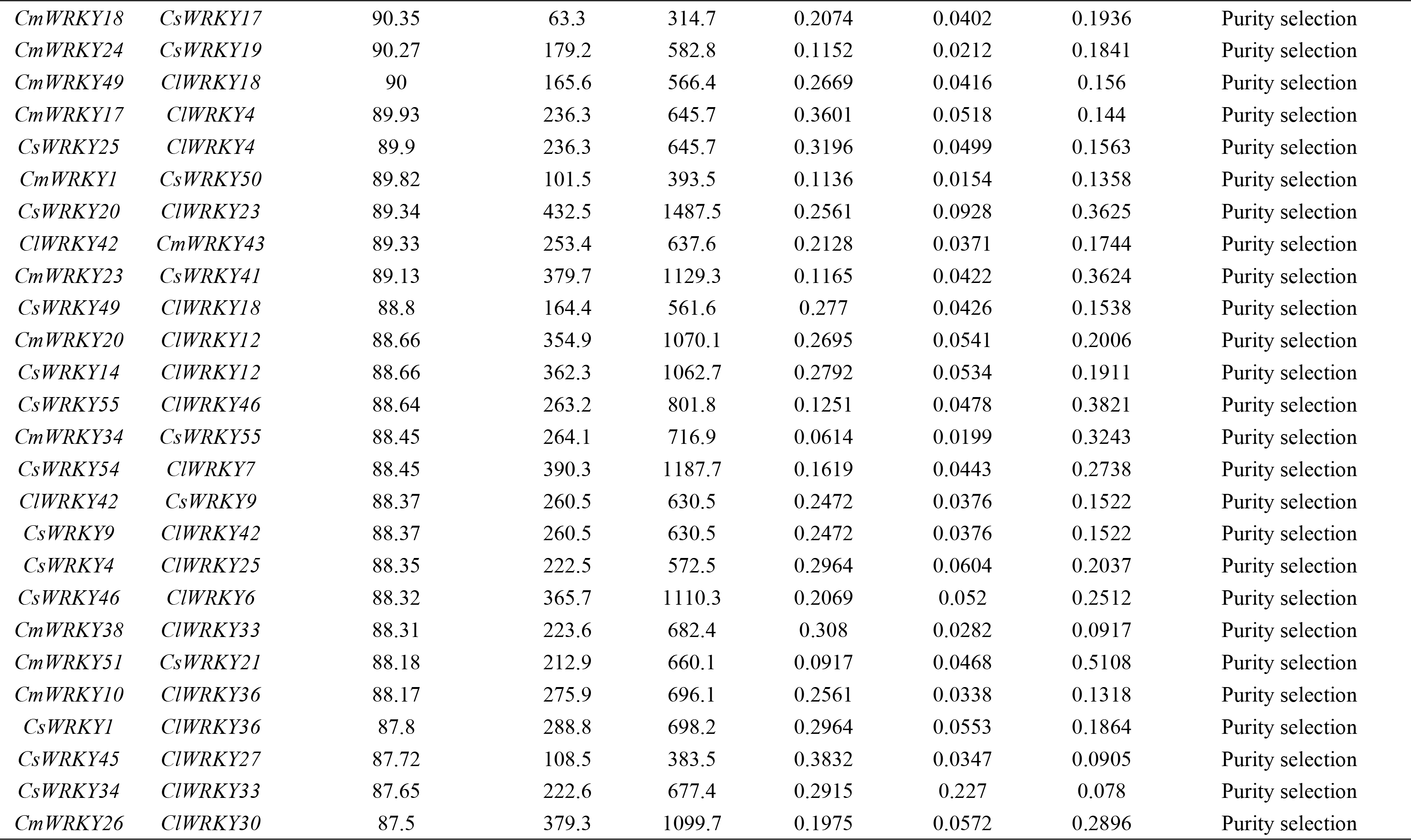

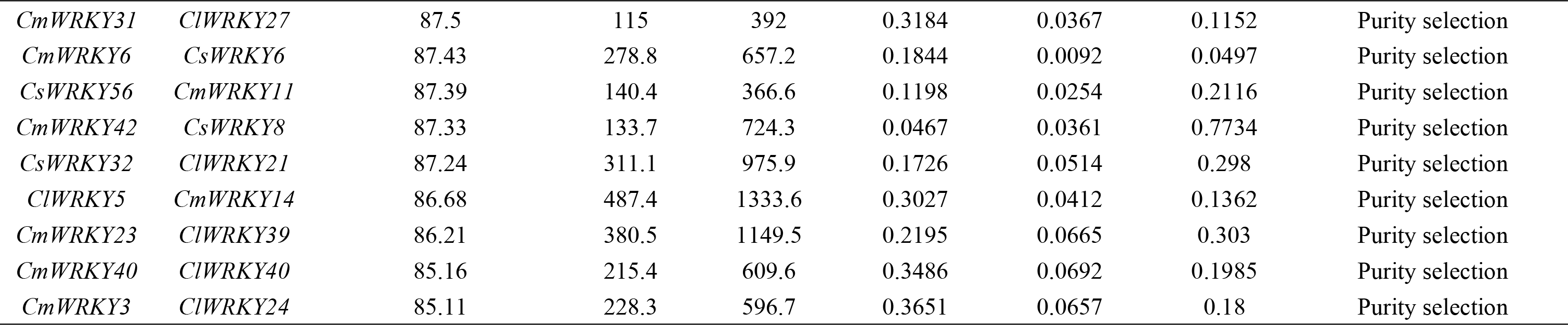

**Figure 4.**
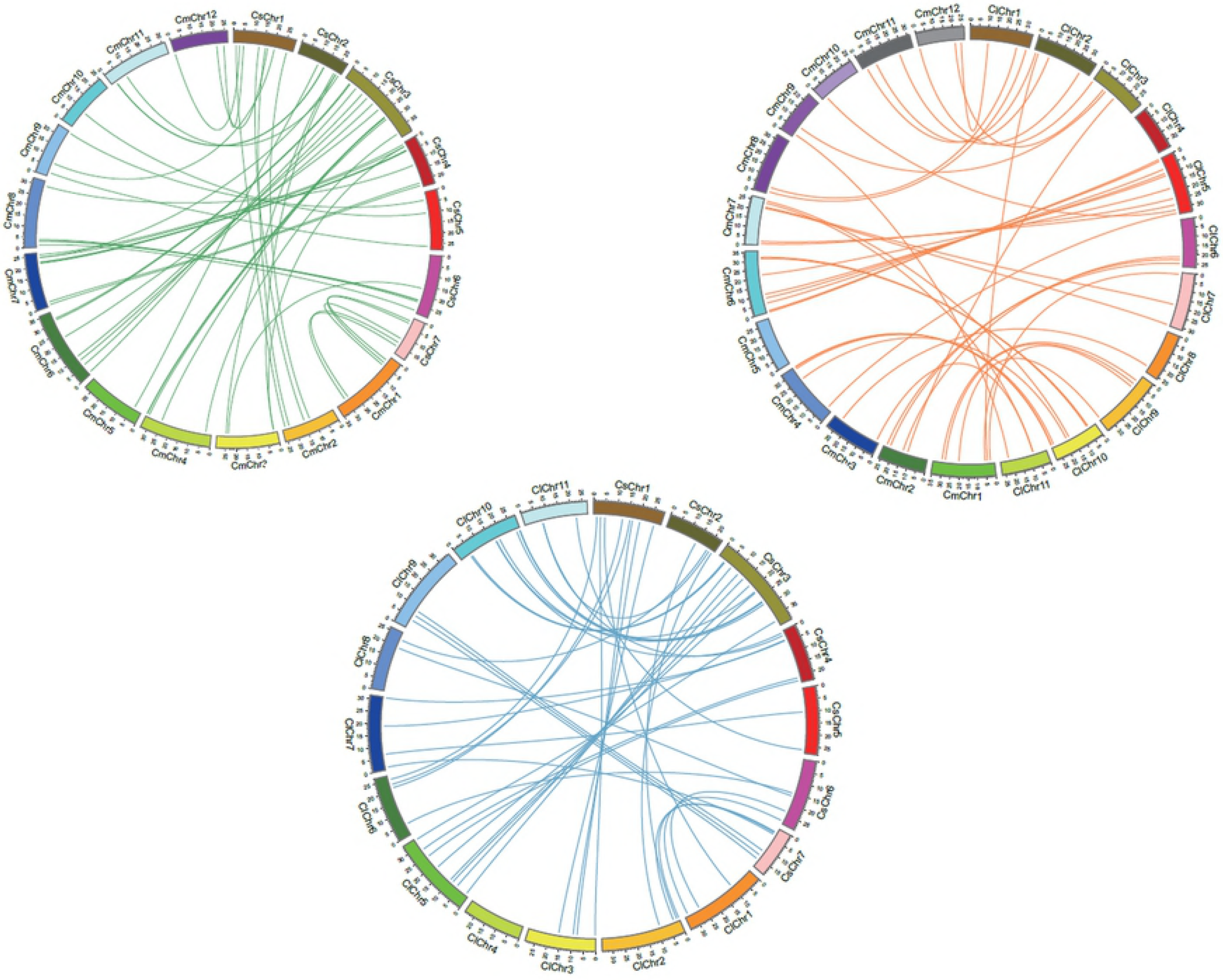
Comparative analysis of orthologous relationship among three *cucurbitaceae* species for *WRKY* genes. Each chromosome was represented with different colour. Green lines indicate homologous genes between melon and cucumber chromosomes, orange lines indicate homologous genes between melon and watermelon chromosomes, blue lines indicate homologous genes between cucumber and watermelon chromosomes.

### Expression patterns of *CmWRKYs* in different tissues at different developmental stages

To gain insights into the potential functions of *CmWRKYs*, the expression patterns of the *CmWRKYs* were investigated using publicly available data from RNA-seq of melon transcript expression generated from 29 different tissues as well as different development stages of melon. The results showed that the transcript abundances of different *CmWRKYs* were extremely diverse (Fig 5). Among the 56 *CmWRKYs*, 27 *CmWRKYs* were detected in all 29 tissues. 55 *CmWRKYs* expect for *CmWRKY5* expressed in root with the transcripts of *CmWRKY6, CmWRKY23, CmWRKY29, CmWRKY40, CmWRKY43, CmWRKY47, CmWRKY51, CmWRKY52, CmWRKY53, CmWRKY55* were more abundant than the other *CmWRKYs*. 51 *CmWRKYs* espressed in leaf with *CmWRKY6, CmWRKY23, CmWRKY28, CmWRKY29, CmWRKY46* and *CmWRKY50* had a high expression levels both in young and old leaves. While *CmWRKY8, CmWRKY17, CmWRKY19, CmWRKY30, CmWRKY49* showed higher expression levels in young leaf than old leaves, whereas *CmWRKY14, CmWRKY16, CmWRKY24, CmWRKY26, CmWRKY27, CmWRKY31, CmWRKY33, CmWRKY34, CmWRKY42, CmWRKY43, CmWRKY47, CmWRKY48, CmWRKY52* and *CmWRKY55* had a higher expression levels in 6 th, 9 th, 12 th leaves than that in young leaf. 46 *CmWRKYs* were expressed in stem with 15 *CmWRKYs* had a higher expression levels. 48 *CmWRKYs* were expressed in tendril. 49 *CmWRKYs* were expressed in flower and ovary, and 50 *CmWRKYs* were expressed in fruit flesh and epicarp. Interestingly, *CmWRKY14, CmWRKY17, CmWRKY20, CmWRKY35, CmWRKY41, CmWRKY43, CmWRKY55* had a higher expression levels in epicarp than that in flesh. Besides, 41 *CmWRKYs* were detected in dry seed, and *CmWRKY19, CmWRKY23, CmWRKY51, CmWRKY52* had a higher expression levels both in flesh and epicarp.

**Figure 5.**
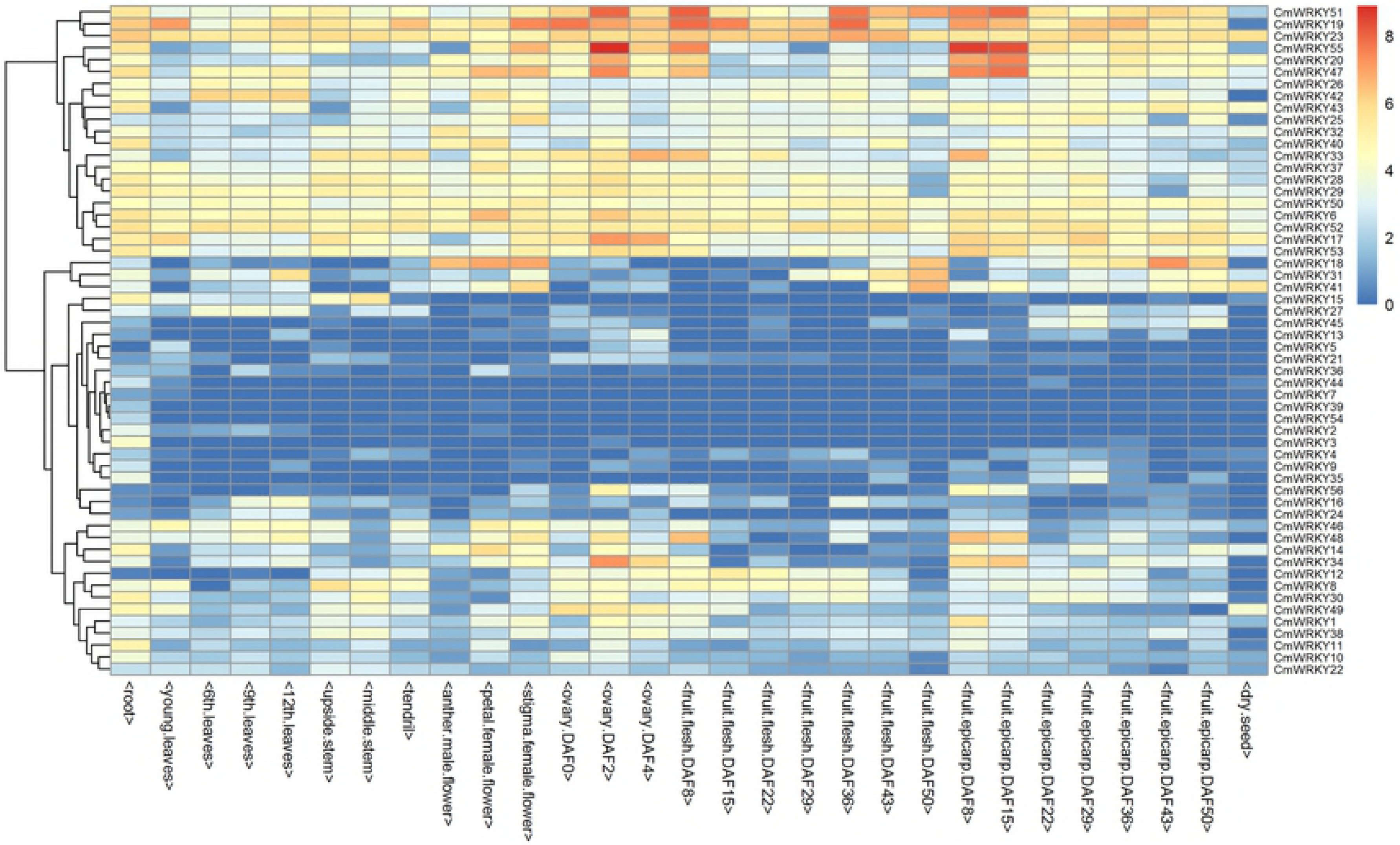
Expression profiles of 56 *CmWRKY* genes in various organs/tissues as well as different developmental stages of melon. The expression values were measured as reads per kilobase of exon model per million mapped reads (RPKM) and shown as log2 (value + 1). The color scale is shown at the right and higher expression levels are shown in red.

### Co-expression analysis of *CmWRKY* and *CmVQ* and in response to powdery mildew infection

It has been previously demonstrated that WRKY proteins can interact physiologically with VQ motif-containing proteins that are involved in the regulation of plant defense responses. In order to gain information about hypothetical interactions between WRKY proteins and VQ proteins in melon, we performed the expression correlation analysis between *CmWRKYs* and *CmVQs* using the data from RNA-seq. Firstly, 24 VQ motif-containing proteins were identified in melon genome. A total of 24 *CmWRKY* genes were co-expressed with 11 *CmVQ* genes with the correlation coefficient was greater than 0.7 (Fig 6; S5 Table). Nine *CmWRKY* genes were simultaneously co-expressed with two different *CmVQ* genes and 15 *CmWRKY* genes were only co-expressed with one *CmVQ* gene. Besides, some *CmWRKY (CmVQ)* genes showed co-expression patterns with other members of *WRKY (VQ)* family genes. Therefore, the co-expression may be important for further functional analysis of *CmWRKYs*.

**Figure 6.**
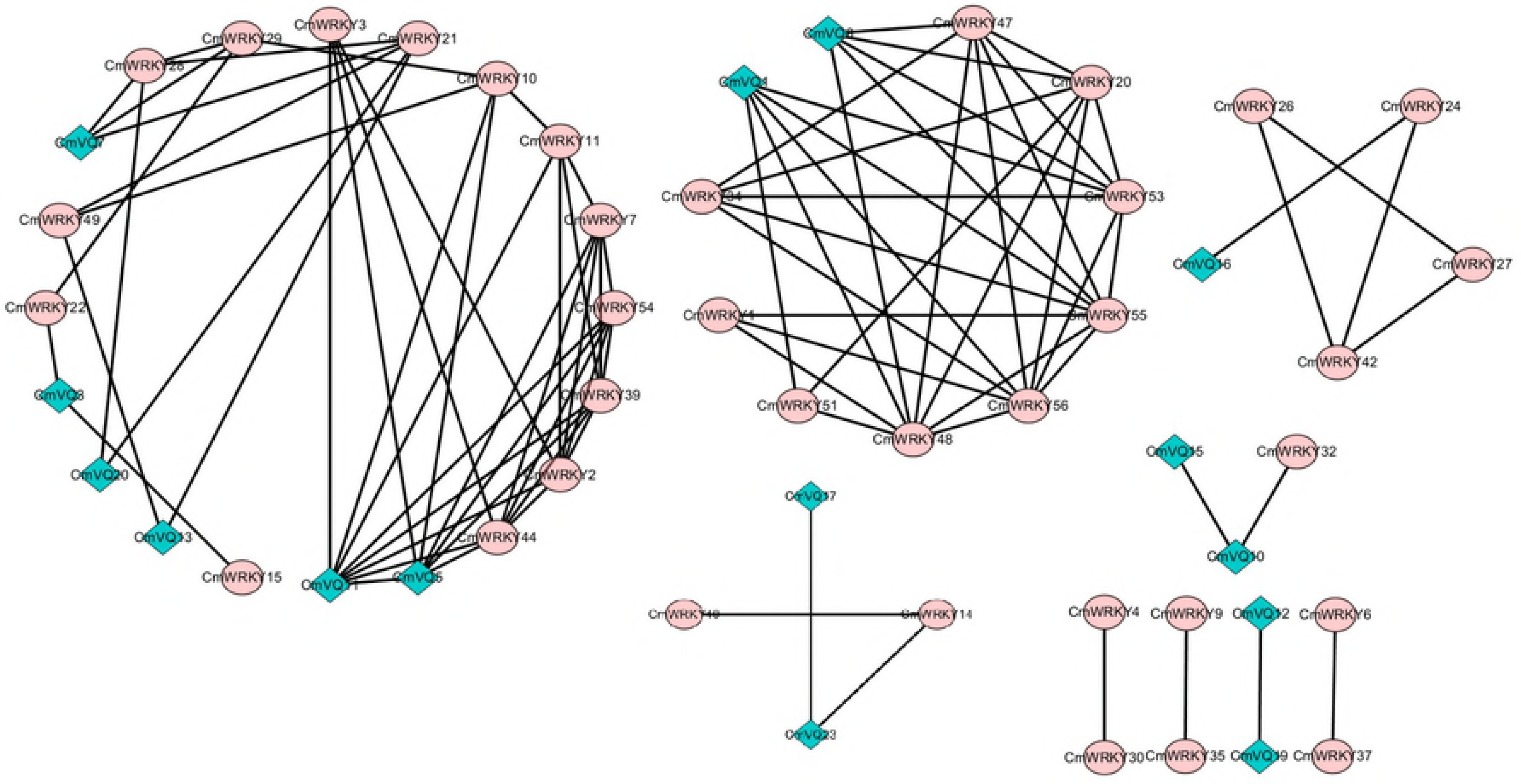
Co-expression network of melon *WRKY* genes and *VQ* genes according to correlation coefficient > 0.7 and p < 0.05. The circles represent *CmWRKY* genes and the diamonds represent *CmVQ* genes.

Numerous reports have indicated that many *WRKY* family genes interact with *VQ* family genes and play important roles in resisting to diverse biotic stresses. To access the function of *WRKY* and *VQ* family genes in resisting to powdery mildew fungus infection, expression profiles of all *CmWRKY* and *CmVQ* in leaves artificially inoculated with powdery mildew fungus were performed based on data from RNA-seq. A total of 16 *CmWRKY* and five *CmVQ* exhibited different expression levels in response to powdery mildew inoculation, indicating that they are powdery mildew fungus responsive genes and might play important roles in resisting to powdery mildew disease in melon (Fig 7). Among them, most *CmWRKY* were up-regulated after inoculation except for *CmWRKY15*. Besides, up-regulation was also observed for *CmVQ6, CmVQ16, CmVQ23*, while *CmVQ18* and *CmVQ21* showed the opposite pattern after inoculation. To validate the RNA-seq data, qRT-PCR was performed to examine the expression of several *CmWRKYs* and *CmVQs* that may be involved in resisting to powdery mildew disease and the results were in agreement with the sequencing data (Fig 8).

**Figure 7.**
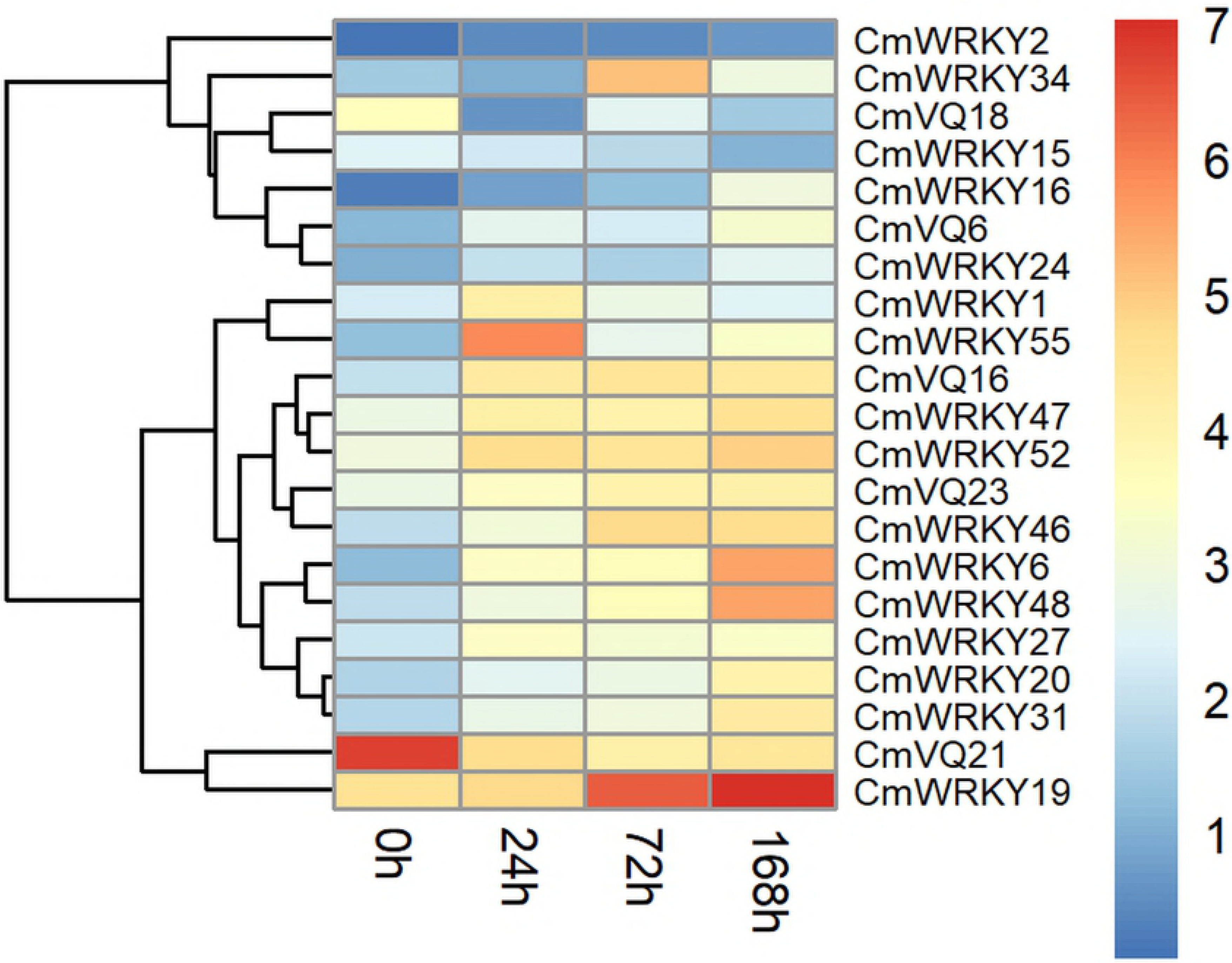
Expression patterns of powdery mildew fungus responsive *CmWRKY* and *CmVQ* genes after powdery mildew fungus infection. The expression values were measured as reads per kilobase of exon model per million mapped reads (RPKM) and shown as log2 (value + 1). The color scale is shown at the right and higher expression levels are shown in red.

**Figure 8.**
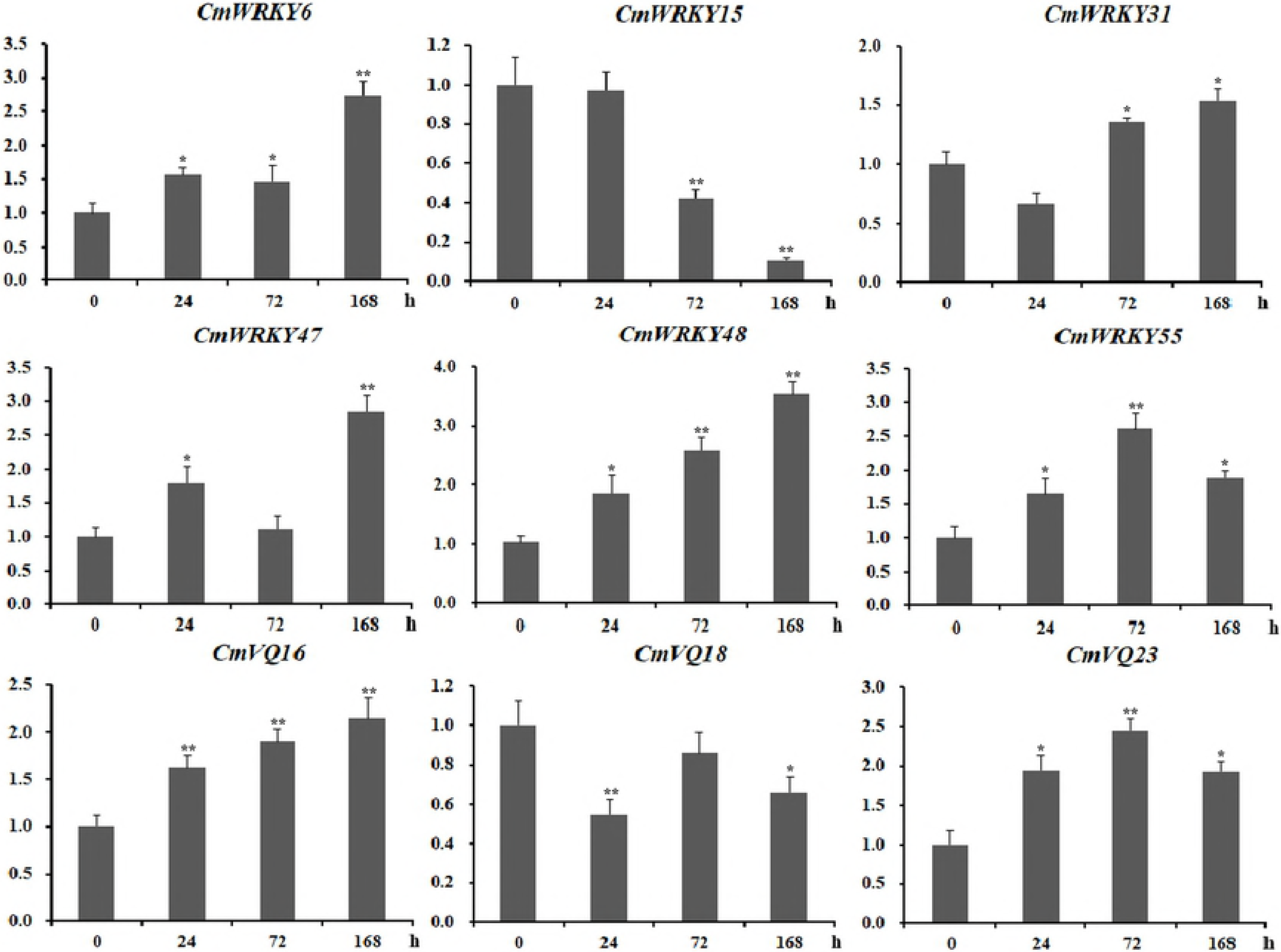
qRT-PCR validation of the differential expression of 9 genes involved in resisting to powdery mildew infection. *CmActin* was used as the internal control. The Y-axis indicates the relative expression level; X-axis (0, 24, 72, 168 h) indicated hours after powdery mildew fungus infection. Error bars indicate the SD of three biological replicates. Asterisks indicated a significant change in expression levels (student’s t-test; *P < 0.05 and **P < 0.01).

## Discussion

### Genome-wide exploration and phylogenetic analysis of *WRKY* genes among cucurbitaceae species

The evolutionary relationship and function analysis of *WRKY* genes have been thoroughly studied in most plants. In the previous report, genome-wide analysis of *WRKY* family genes in cucumber and watermelon has also performed, and 55 *CsWRKYs* as well as 63 *ClWRKYs* in the cucumber and watermelon genome were identified, respectively [11, 14]. To further gain information with respect to the evolutionary relationship and function of *WRKY* genes among *cucurbitaceae* species, we exploited available genomic resources to characterize the *WRKY* family genes in the other *cucurbitaceae* species, melon. A total of 56 putative *WRKY* genes (proteins) were identified in the genome of melon. The numbers of *WRKY* family members in melon were approximately equal to that in cucumber (56) and grape (55) but less than that in *Arabidopsis* (72 members) and in rice (102 members), peanut (152) and other crops, which maybe because unlike *Arabidopsis* and rice, *cucurbitaceae* species and grape have a much smaller genome and have not undergone recent whole-genome duplication after the differentiation of eudicots and monocots. Indeed, previous studies showed that *Arabidopsis* has undergone three whole-genome duplication events, and rice has also undergone at least one whole-genome duplication event, which promoted the rapid expansion of gene family [35, 36]. However, Huang and Garcia-Mas have respectively observed the absence of recent whole-genome duplications in cucumber and melon genomes [28, 29].

It is worth noting that ClWRKY11 that contain one WRKY domain were clustered in group IN from the phylogenetic analusis. Meanwhile, two CsWRKY, CsWRKY46 and CsWRKY51, were clustered in group IN, and CsWRKY47 and CsWRKY52 were clustered in group IC. Thus, these WRKYs in group II may have arisen from a two-domain WRKY protein in group I that lost one of its WRKY domains located in the N-terminal or C-terminal during evolution. The previous studies have reported that group I, group II, group III contained 13, 45, 14 WRKY proteins in *Arabidopsis* and 13, 42, 47 WRKY proteins in rice, respectively [37]. Compared with the numbers of *WRKY* families in *Arabidopsis* and rice, it is apparent that variations in the number of *WRKY* genes in group III are the primary cause of the diversity of *WRKY* gene family size between melon and other plants. Therefore, a key role of group III *WRKY* genes in plant evolution may exist, which is possible that the genome or gene family duplication events have resulted in the different size of the group III *WRKY* genes among *cucurbitaceae* species, *Arabidopsis* and rice.

### The orthologous and expression analysis of *WRKY* genes provide important clues for their function

The recent gene duplication events including segmental duplication and tandem duplication are most important in the expansion and evolution of gene families, and were considered to be the raw materials for new biological functions. Therefore, we further analyzed the influence of recent duplication events to *WRKY* family genes in *cucurbitaceae* species. The results showed that there are no recent tandem duplication and segmental duplication event in *WRKY* family genes of *cucurbitaceae* species, as the nucleotide identity of paralogous pairs were lower than 80%. Therefore, the absence of recent duplication events in melon, watermelon and cucumber genome may attribute to the small size of *WRKY* members.

Given that orthologous genes among different plants are generally supposed to retain similar functions and to share other key properties. Thus, the comparative analysis of *WRKY* orthologous genes among *cucurbitaceae* species could help to predict their genetic relationship and the potential functions of WRKY proteins in melon, cucumber and watermelon. For example, *AtWRKY46* played very important roles in response to drought and salt stresses in *Arabidopsis*, and as the orthologous gene of *AtWRKY46* in watermelon, *ClWRKY23*, was also reported to be up-regulated under drought and salt stresses, implying that these two *WRKY* play similar functions in different plant species [14]. In our study, a total of 44 orthologous pairs between melon and watermelon, 43 orthologous pairs between melon and cucumber were identified, which provided important clues for further functional prediction of *WRKY* genes in melon. A WRKY transcription factor from cucumber, *CsWRKY46*, confers cold resistance in transgenic plants by regulating a set of cold-stress responsive genes in an ABA-dependent manner [38]. As its orthologous genes, *CmWRKY29* may also play similar roles in response to cold stress in melon. However, further molecular and biological experiments should be carried out to investigate their biological function.

WRKY transcription factors play very critical roles in different tissues or different developmental stages to control plant growth and development. For instance, virus-induced silencing of *GmWRKY58* and *GmWRKY76* in soybean causes severe stunted growth with reduced leaf size and plant stature, and overexpression of the *OsWRKY31* could reduce lateral root formation and elongation [39, 40]. To provide important clues for gene function prediction, thus, we conducted a digital gene expression analysis for *CmWRKY* genes in different tissues including leaf, root, stem, tendril and flower ovary and fruit as well as their different developmental stages using released data from RNA-seq. The temporal and spatial diversification of *WRKY* gene expression is widespread, which is important for gene function analysis. *CmWRKY6, CmWRKY17, CmWRKY19, CmWRKY23, CmWRKY42, CmWRKY47* and *CmWRKY52* had a higher expression level in all tested tissues, implying that these genes play key roles in the whole-plant growth and development. Furthermore, many *CmWRKYs* had a higher expression level in certain organs/tissues or in different developmental stages, which suggested that they play divergent roles in the different developmental processes.

### CmWRKY might play important roles in resistance to powdery mildew disease through interaction with CmVQ

In addition to the roles of WRKY proteins in plant growth and development, increasing reports indicated that WRKY proteins in various plant species are involved in the response to various biotic and abiotic stresses. Moreover, many WRKY proteins are also the components of plant biotic stress regulatory networks and directly regulate the expression of several critical genes of defense-signaling pathways [41, 42]. For example, *AtWRKY33* can positively modulate defence-related gene expression and improve resistance to *B. cinerea* disease, while *AtWRKY7* and *AtWRKY48* have an immediate negative effect on plant defense response [43–47]. Overexpression of *OsWRKY67* in rice confirmed enhanced disease resistance to *Magnaporthe oryzae* and *Xanthomonas oryzae*, but led to a restriction of plant growth in transgenic lines [48]. Furthermore, Guo demonstrated that 16 *VvWRKY* have a negative effect on grape powdery mildew resistance and 22 *VvWRKY* have a positive effect on grape powdery mildew resistance in grape [49]. Transcriptome analysis and qRT-PCR expression profiles in melon leaves generated in the current study also revealed that many WRKY genes were responsive to powdery mildew inoculation, thus highlighting the extensive involvement of *WRKY* genes in resisting to powdery mildew disease.

Previous reports showed that some rice and grape *WRKY* genes were co-expressed with *VQ* genes during the response to attacks by three different pathogens [50, 51]. Further studies have revealed that WRKY proteins can interact physiologically with VQ motif-containing proteins [2, 52]. Thus, VQ and WRKY proteins might assemble to form one complex to regulate the target gene. Indeed, VQ9 protein acts as a repressor of the WRKY8 protein to modulate salinity stress tolerance and pathogen resistance in *Arabidopsis* [53]. In the present study, we found that 24 *CmWRKY* genes were co-expressed with 11 *CmVQ* transcription factors (Fig 6). Furthermore, some of the co-expressed *CmWRKY* and *CmVQ* genes were shown to be response to powdery mildew infection, implying that these VQ and WRKY proteins are involved in the same biological pathway. For example, *CmVQ6* positively co-expressed with *CmWRKY47, CmWRKY55, CmWRKY56* and all these were up-regulated by powdery mildew inoculation (Fig 6 and 7), suggesting that different WRKY have the same VQ partner. From above results, we have shown which WRKY proteins interact with which VQ proteins. Further research should carried out to explore their physical interactions in vitro during the responses to powdery mildew infection as well as various abiotic stresses in melon to provide further molecular evidence for these interactions.

## Conclusions

In this study, we identified 56 WRKY family genes in melon. WRKY proteins in *cucurbitaceae* species were under strong purifying pressure. The expression pattern analysis of *CmWRKYs* in different tissues as well as under powdery mildew infection indicated that they were involved in the growth and development of various tissues and that they might positively or negatively participated in plant tolerance against powdery mildew disease. Furthermore, the co-expression analysis between *Cm WRKY* and *CmVQ* will assist in understanding the roles of these *CmWRKY* transcription factors in response to powdery mildew disease and their potential interactions among defence-related genes in the disease resistance network. Collectively, our findings provide valuable clues for further research on the specific function and regulatory mechanisms of *WRKY*s in *cucurbitaceae* species, and could help to select appropriate candidate genes for further characterization of their pathogen resistant functions in melon.

## Author contributions

CG designed the research and performed the bioinformatics analysis. JLS contributed to RNA extraction and qRT-PCR. ZGJ wrote the original manuscript. YMD finalized the figures and tables. CQW and SHX managed the plant materials and analyzed the data. XLG and LBL reviewed the grammar. QWC and WDL revised the original manuscript. All authors have read and approved the final manuscript.

## Acknowledgements

This work was financially supported by grants from the National Technical System of Watermelon and Melon Industry (CARS-25), the Natural Science Foundation of Shandong Province (ZR2018LC017) and Agricultural Science and Technology Innovation Project of Shandong Academy of Agricultural Sciences (CXGC2018E08).

## Supporting Information

**Table S1.** Oligonucleotide primer sequences used for qRT-PCR.

**Table S2.** The full-length nucleotide sequences and amino acid sequences of all identified WRKY family members in three *cucurbitaceae* species.

**Table S3.** The detailed information of all identified WRKY family members in melon.

**Table S4.** The information of orthologous gene pairs among *cucurbitaceae* species.

**Table S5.** The correlation coefficient values of *CmWRKY* and *CmVQ* genes.

